# Female grant applicants are equally successful when peer reviewers assess the science, but not when they assess the scientist

**DOI:** 10.1101/232868

**Authors:** Holly O. Witteman, Michael Hendricks, Sharon Straus, Cara Tannenbaum

## Abstract

**Background:** Previous research shows that men often receive more research funding than women, but does not provide empirical evidence as to why this occurs. In 2014, the Canadian Institutes of Health Research (CIHR) created a natural experiment by dividing all investigator-initiated funding into two new grant programs: one with and one without an explicit review focus on the caliber of the principal investigator.

**Methods:** We analyzed application success among 23,918 grant applications from 7,093 unique principal investigators in a 5-year natural experiment across all investigator-initiated CIHR grant programs in 2011-2016. We used Generalized Estimating Equations to account for multiple applications by the same applicant and an interaction term between each principal investigator’s self-reported sex and grant programs to compare success rates between male and female applicants under different review criteria.

**Results:** The overall grant success rate across all competitions was 15.8%. After adjusting for age and research domain, the predicted probability of funding success in traditional programs was 0.9 percentage points higher for male than for female principal investigators (OR 0.934, 95% CI 0.854-1.022). In the new program focused on the proposed science, the gap was 0.9 percentage points in favour of male principal investigators (OR 0.998, 95% CI 0.794-1.229). In the new program with an explicit review focus on the caliber of the principal investigator, the gap was 4.0 percentage points in favour of male principal investigators (OR 0.705, 95% CI 0.519- 0.960).

**Interpretation:** This study suggests gender gaps in grant funding are attributable to less favourable assessments of women as principal investigators, not differences in assessments of the quality of science led by women. We propose ways for funders to avoid allowing gender bias to influence research funding.

**Funding:** This study was unfunded.

## INTRODUCTION

For decades, studies have shown that women in academia and science must perform to a higher standard than men to receive equivalent recognition,^1–4^ especially Indigenous and racialized women.^5–11^ Compared to men, women are more often characterized as lacking the brilliance, drive, and talent required to carry a novel line of inquiry through to discovery,^8,12^ with children as young as six years old endorsing such stereotypes.^13^ Women are less likely than men to be viewed as scientific leaders^14–17^ or depicted as scientists.^18^ Women in academia contribute more labor for less credit on publications,^19,20^ receive less compelling letters of recommendation,^21–24^ receive systematically lower teaching evaluations despite no differences in teaching effectiveness,^25^ are more likely to experience harassment,^11,26–29^ and are expected to do more service work^30,31^ and special favors for students.^32^ While men in academia have more successful careers after taking parental leave, women’s careers suffer after the same.^33^ Women receive less start-up funding as biomedical scientists^34^ and are underrepresented in invitations to referee papers.^35^ Compared to publications led by men, those led by women take longer to publish^36^ and are cited less often,^37,38^ even when published in higher-impact journals.^39^ Papers^40^ and conference abstracts^41^ led by women are accepted more frequently when reviewers are blinded to the identities of the authors. Women are underrepresented as invited speakers at major conferences,^4^ prestigious universities,^42^ and grand rounds.^43,44^ When women are invited to give these prestigious talks, they are less likely to be introduced with their formal title of Doctor.^45^ Female surgeons have been shown to have better patient outcomes overall,^46^ yet, when a patient dies in surgery under the care of a female surgeon, general practitioners reduce referrals to her and to other female surgeons in her specialty, whereas they show no such reduced referrals to male surgeons following a patient’s death.^47^ Women are less likely to reach higher ranks in medical schools even after accounting for age, experience, specialty, and measures of research productivity.^48,49^ When fictitious or real people are presented as women in randomized experiments, they receive lower ratings of competence from scientists,^50^ worse teaching evaluations from students,^51^ and fewer email responses from professors after presenting as students seeking a PhD advisor^9^ or as scientists seeking copies of a paper or data for a meta-analysis.^52^

Conversely, other research has demonstrated advantages experienced by women in academia; for example, achieving tenure with fewer publications than men.^53^ In assessments of potential secondary and postsecondary teachers and professors, women are favored in male-dominated fields, as are men in female-dominated fields.^54^ When fictitious people are presented as women 2 in randomized experiments, they receive higher rankings as potential science faculty.^55,56^ This aligns with evidence from other contexts showing that high-potential women are favored over high-potential men^57^ and that, while women face discrimination at earlier stages, once women have proven themselves in a male-dominated context, they are favored over men.^58^

In sum, there is considerable evidence that women face or have faced persistent, pervasive barriers in many aspects of academia and science. There is also recent evidence that women in academia and science may fare better in merit evaluations than equally-qualified men after they have progressed beyond postsecondary education. In light of this evidence as a whole, we consider the question: does bias in favor of men or women influence research grant funding?

Previous research on this question has been suggestive but not conclusive. A 2007 meta-analysis of 21 studies from a range of countries found an overall gender gap in favor of men, with 7% higher odds of fellowship or grant funding for male applicants.^59^ Research since then has documented that, compared to their male colleagues, female principal investigators have lower grant success rates,^60^ lower grant success rates in some but not all programs,^61,62^ equivalent grant success rates after adjusting for academic rank^63,64^ but fewer funds requested and received,^63–66^ or equivalent funding rates.^67^ To the best of our knowledge, no such study has yet found women to experience higher grant success rates nor to receive more grant funding than men.

These previous studies have been observational, making it difficult to draw robust conclusions about the causes of sex-related discrepancies in funding success rates when they are observed. Furthermore, some previous studies have not accounted for potential confounding variables; for example, domain of research.^59,68,69^ Our objective in this study was therefore to determine whether gender gaps in grant funding are attributable to potential differences in how male and female principal investigators are evaluated, using real-world data and a study design that would allow for stronger conclusions than those from observational studies. Our study made use of a natural experiment at a national health research funding agency which allowed for the comparison of grant success rates among male and female principal investigators between three grant programs. The three programs were: traditional grant programs, which had demonstrated higher success rates among younger male principal investigators for applications submitted in 2001-2011^60^ but for which no such analysis had been conducted after 2011, and 3 two new competitions, one with and without an explicit review focus on the quality of the principal investigator.

Our hypotheses regarding comparisons of gaps in grant success rates between female and male principal investigators after accounting for age and research domain were as follows.

> H_0_: Gaps will be similar under traditional review criteria and both sets of new review criteria.
>
> H_1_: The gap will be larger in favour of male principal investigators in the new competition with more focus on the science.
>
> H_2_: The gap will be larger in favour of male principal investigators in the new competition with more focus on the scientist.

Support for H_0_ would suggest that gaps, when present, may reflect different career paths and choices made by women and men,^70,71^ differences between the types of research proposed by female and male principal investigators, or may be spurious. Support for H_1_ would suggest that gaps are are partly or wholly driven by female principal investigators proposing science assessed as lower quality than that of their male colleagues. Support for H_2_ would suggest that gaps are partly or wholly driven by women being assessed less favorably as principal investigators compared to their male colleagues. Other potential alternative hypotheses such as gaps in favour of female principal investigators were not considered a priori because publicly-available summary statistics of the programs showed such results to be impossible.

## METHODS

### Population, Setting and Data Source

As depicted in Figure 1, beginning in 2014, the Canadian Institutes of Health Research (CIHR) phased out traditional open grant programs, dividing investigator-initiated funding into two new open programs: the Project grant program and Foundation grant program. Both new programs used a staged review process in which lower-ranked applications are rejected from continuing on to further stages. As in traditional programs, in the new Project grant program, reviewers were instructed to primarily assess the research proposed. In contrast, the Foundation grant program was about ‘people, not projects’. Reviewers at the first stage were instructed to primarily assess the principal investigator, with 75% of the score being allocated to reviewers’ assessments of the applicant’s leadership, productivity, and the significance of their 4 contributions.^72^ Table 1 lists the adjudication criteria for each of these. Only applicants who passed this stage were invited to submit a detailed proposal describing their research.

**Table 1.** Foundation Grant Stage 1 Application Adjudication Criteria

| Application Section | Maximum Length | Score Weight | Criteria |
| --- | --- | --- | --- |
| Leadership | 1750 characters, including spaces (about half a page) | 25% | • Is the applicant recognized in their field, demonstrating a history of holding influential roles, inspiring others, mobilizing communities and advancing the direction of a field?<br>• Has the applicant demonstrated the ability to successfully establish, resource, and direct programs of research, which should include: securing the required resources, ensuring effective collaboration, and/or incorporating knowledge translation strategies? |
| Significance of Contributions | 1750 characters, including spaces (about half a page) | 25% | • Has the applicant significantly advanced knowledge and/or its translation into improved health care, health systems, and/or health outcomes?<br>• Has the applicant engaged, trained, and/or launched the career paths of promising individuals in research and/or other health-related non-academic fields? |
| Productivity | 1750 characters, including spaces (about half a page) | 25% | • Has the applicant demonstrated an outstanding level (quantity) of research outputs in their field based on prior work?<br>• Has the applicant's previous work generated high quality research outputs? |
| Vision and Program Direction | 3500 characters, including spaces (about a page) | 25% | • Are the vision, goal, overall objective(s), and potential contribution(s) of the proposed research program well-defined and well-articulated in the context of a logical career progression for the Program Leader(s)?<br>• Is the vision forward-looking, creative, and appropriately ambitious?<br>• Does the vision aim to significantly advance knowledge and/or its translation to improved health care, health systems, and/or health outcomes? |

**Figure 1.**
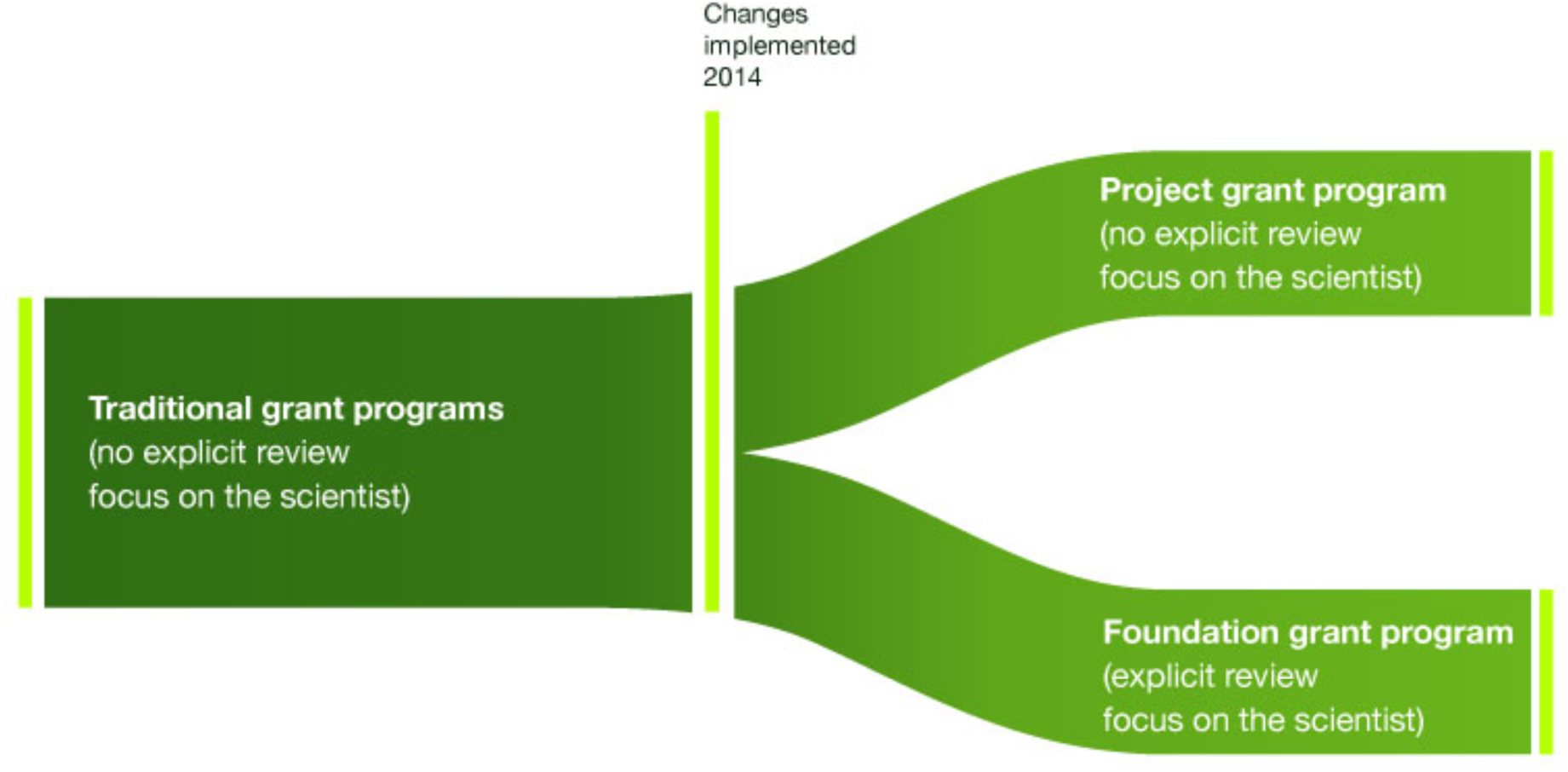
Changes in grant programs.

### Analysis

We analyzed data from all applications submitted to CIHR grant programs across all investigator-initiated competitions from 2011 through 2016. We used Generalized Estimating Equations to fit a logistic model that accounted for applicants submitting multiple applications during these five years.^73,74^ We modeled grant success rates as a function of the grant program, applicants’ self-reported binary sex, self-reported age, self-declared domain of research, and an interaction term between applicant sex and grant program. We controlled for age and domain of research because younger cohorts of investigators include larger proportions of women, as do domains of health research other than biomedical. Full methodological details are available in the online appendix.

## RESULTS

The data set contained 23,918 applications from 7,093 unique applicants. There were 15,775 applications from 4,472 male applicants and 8,143 applications from 2,621 female applicants. The overall grant success rate across the data set was 15.8%. As shown in Figure 2, after adjusting for age and research domain, the predicted probability of funding was 0.9 percentage points higher for male applicants than female applicants in traditional grant programs (OR 0.934, 95% CI 0.854-1.022), 0.9 percentage points higher in the Project program (OR 0.998, 95% CI 0.794-1.229) and 4.0 percentage points higher in the Foundation program (OR 0.705, 95% CI 0.519-0.960). Figure 3 shows how the gap in the Foundation program was driven by discrepancies at the first review stage, where review focused on the principal investigator. Odds of receiving funding were also lower in the three non-biomedical domains and higher for older applicants. Tabular results are available in the online appendix.

**Figure 2.**
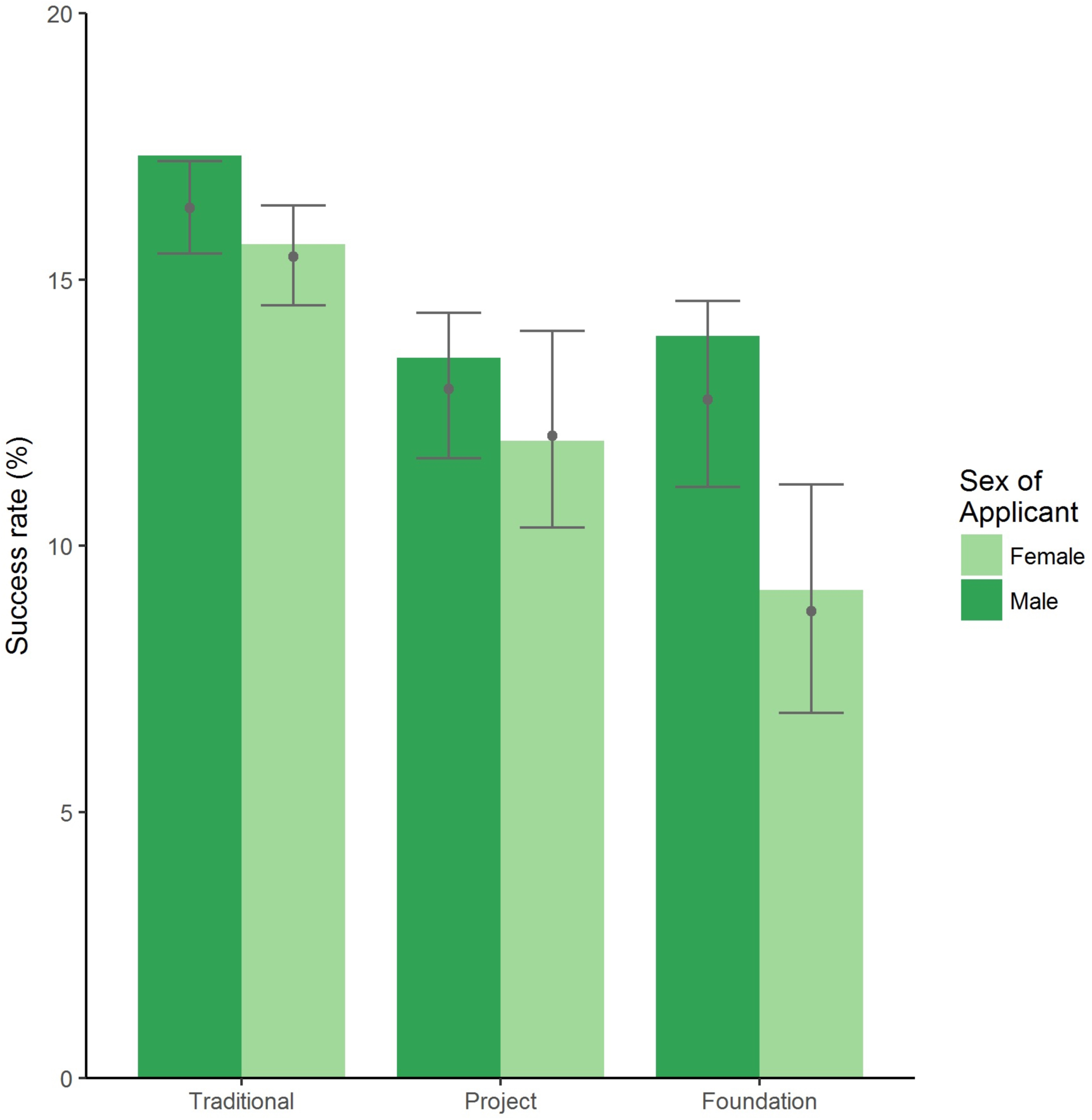
Funding success rate by grant program. Columns indicate observed success rates. Points and error bars indicate model-predicted means and 95% confidence intervals, respectively.

**Figure 3.**
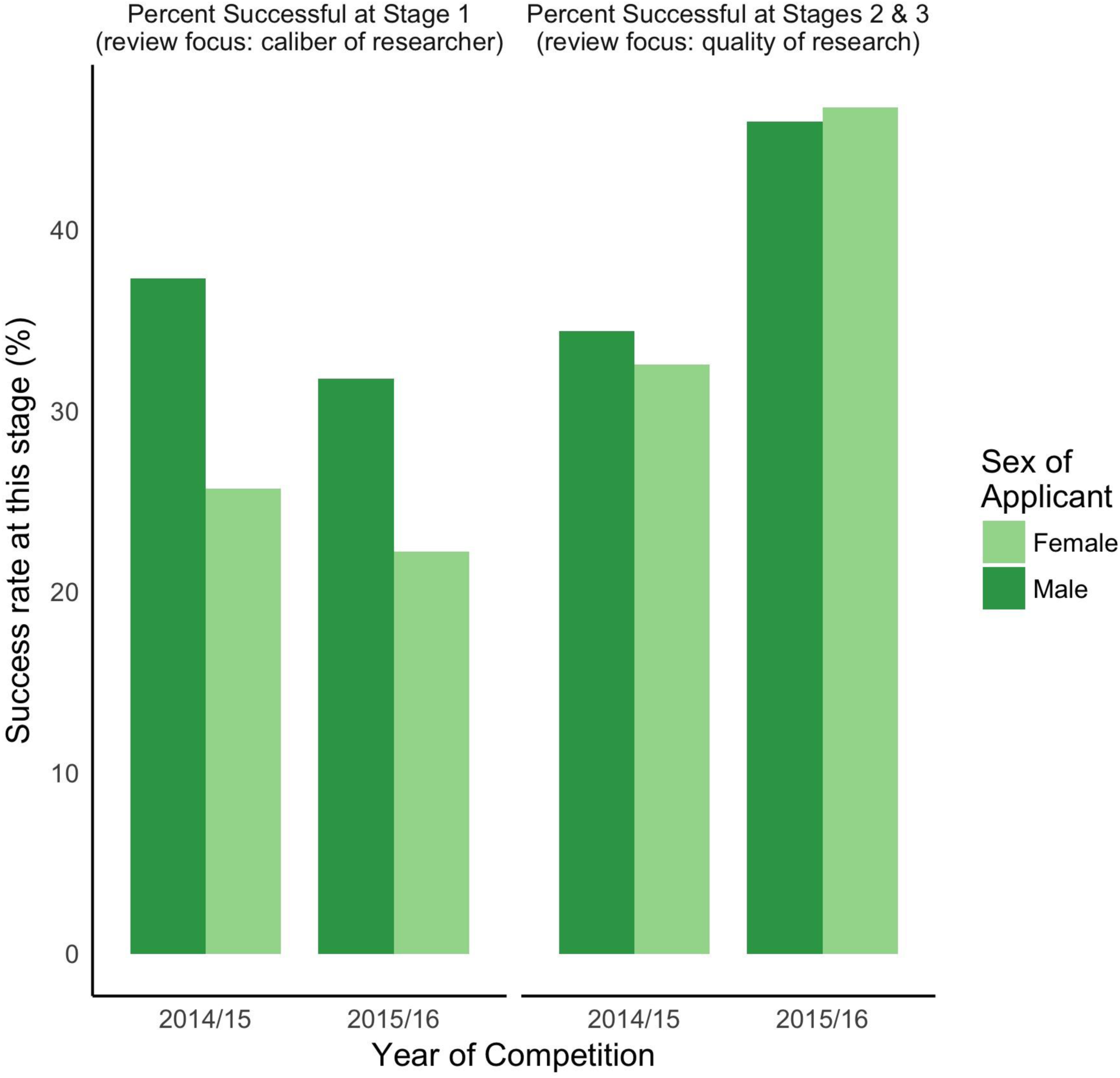
Foundation results by review stage. Columns indicate observed success rates.

## DISCUSSION

Our study provides stronger evidence than was previously available regarding the likely causes of gender gaps in grant funding. In this natural experiment, when reviewers primarily assessed the science, there were no statistically significant differences between success rates for male and female principal investigators. When reviewers explicitly assessed the principal investigator as a scientist, the gap was significantly larger. These data support the second of our alternative 5 hypotheses; namely, that gender gaps in funding stem from female principal investigators being evaluated less favourably than male principal investigators, not from differences in evaluations of the quality of their science.

The finding that women are evaluated less favourably than men as applicants aligns with previous studies from other countries. Data from the United States showed that female grant applicants to the National Institutes of Health’s flagship R01 program were less likely than male applicants to be described as leaders.^1^ ^4^ In the Netherlands, grant reviewers gave equal scores to men’s and women’s proposed research but assigned lower scores to women as researchers.^75^ In Sweden, similar differences have been shown among evaluators’ assessments of applicants for governmental venture capital.^76^

There are two main reasons that female principal investigators might be evaluated less favourably than male principal investigators. First, reviewers may exhibit gender-biased subjective evaluations of principal investigators. Second, female principal investigators may have submitted weaker applications than male peers. Below, we consider each of these two potential reasons in the context of available literature.

The suggestion that reviewers might exhibit conscious or unconscious bias against women aligns with previous evidence about gender gaps at the highest levels of achievement. For example, gender gaps in top mathematics performers have been shown to be attributable to cultural gender inequality, not differences in natural ability.^77^ In orchestra auditions, women became more successful when auditioning musicians’ identities were concealed behind a screen.^78^ Similar increases in success have occurred for women submitting manuscripts^40^ and conference abstracts^41^ after reviewers were blinded. Randomized experiments using hypothetical scenarios (e.g., “How would you respond if …”) have shown scientists to favour hypothetical men as laboratory managers,^50^ hypothetical men as junior faculty in an experiment conducted in 1997-1998,^79^ but hypothetical women as junior faculty in a more recent experiment.^55,56^ Randomized experiments in which the outcome was real behaviour all favoured men,^51,52^ particularly white men.^9^

If Foundation Grant reviewers’ evaluations of applicants were gender-biased, it is possible that such bias might have been due to elements outside reviewers’ control; for example, the structure of the application. The Foundation Grant Stage 1 application form was accompanied 6 by a structured curriculum vitae (CV) describing the last seven years of the principal investigator’s research. During this study, the seven-year limit was not extended in the case of a maternity, parental, or other leave. It is possible that the inability to list a full seven years’ worth of productivity in the case of leaves may have more negatively influenced reviewers’ evaluations of female compared to male applicants. Female principal investigators may also have described their accomplishments more modestly, though, to our knowledge, there are no data suggesting this gendered stereotype applies to competitive scientists.

The idea that reviewers’ evaluations might reflect weaker applications by female principal investigators requires that female principal investigators showed weaker leadership, less significant contributions, lower productivity, or less compelling visions and program directions compared to male principal investigators. Regarding leadership, previous evidence shows women are less likely to advance to leadership roles in medicine, even after accounting for confounding variables such as differences in publication records.^49,80^ Women’s CVs are less likely to include prestigious invited talks.^4,42–44^ The leadership credentials of women in health research may also be constrained by broader societal tendencies in which women receive negative responses for being ambitious^81^ or exerting control as leaders.^82–84^ Research suggests that women must choose between being seen as a competent leader or being liked,^82,83^ which complicates career advancement within academia, where leadership positions often depend on colleagues’ endorsements. Societal stereotypes suggest that women are less interested in leadership positions, but to our knowledge there is no evidence that this is true for women in health research.

Regarding significance of contributions, papers led by women have been shown to be cited less often,^37,38^ which may indicate lower significance of contributions. It is possible that women’s track records of impactful work may be negatively affected by expectations of service and other work^30–32^ or by the “productivity tax”^85^ of dealing with harassment.^11,26–28^ Data are lacking about men and women’s records regarding launching the careers of trainees, but research suggests that trainees publish more under the supervision of female principal investigators.^86,87^

Regarding productivity, globally, there are more publications authored by men.^37,88^ Ignoring the potential confounder of different publication norms in fields with different gender ratios, papers from Canada had a somewhat lower proportion of female fractional authorships (31% for publications in 2008-2012^89^) than national statistics of female faculty might suggest (35% at 7 rank assistant professor or higher in 2010-2011^90^). Male principal investigators in our study may also have had more publications due to historically-higher CIHR grant funding yielding more data for them to publish.^60^ Together, this evidence suggests that it is possible that female principal investigators in our study had fewer publications compared to their male peers, though this may be confounded by career stage. Research by Tamblyn and colleagues is examining male and female principal investigators’ productivity records within the Foundation grant program and whether these were evaluated equivalently by reviewers. Older research showed that reviewers in Sweden assessed equivalent productivity less positively for female fellowship applicants compared to male applicants,^2^ but this study analyzed reviews conducted nearly twenty-five years ago and may not apply in our context.

Finally, regarding vision and program direction, to the best of our knowledge, there is no previous evidence about gender gaps or lack thereof in evaluations of such brief descriptions of research programs. The lack of difference in evaluations of Project Grants suggests that it is unlikely there were inherent differences in the quality of program descriptions.

In summary, we cannot pinpoint whether female principal investigators may have been evaluated more negatively due to bias on the part of reviewers or to review criteria designed around metrics that may reflect lower performance or cumulative disadvantage. Previous literature suggests that both individual and structural bias are plausible. There is more experimental evidence supporting the potential for individual bias than structural bias or objective differences in performance, but it is arguably harder to run randomized experiments about structural bias. Perhaps most importantly, from a policy perspective, it matters little which of these is the most likely cause. If, indeed, there are systematic differences between the leadership, contribution and productivity metrics of male and female principal investigators, is a funding competition based on these metrics justified when there is no indication that women’s science is of lower quality?

The CIHR deemed such imbalanced outcomes not justifiable. Following the grant cycles analyzed in our study, and as part of a broader Equity Framework,^91^ the CIHR implemented new policies to eliminate the observed gender gap in Foundation grants. The policies included instructions that reviewers should complete an evidence-based^92^ reviewer training module about multiple forms of unconscious bias.^93^ Training has been shown to help mitigate the effects of bias in some studies^15,94^ though not in others.^95,96^ The Foundation grant program regulations 8 were also revised such that, should reviewer training not have the desired effects at Stage 1, a proportional number of female applicants would nonetheless proceed to the next stage at which their proposed research would be evaluated. In other contexts, such quotas based on the available pool of candidates have been shown to increase women’s representation, the overall quality of candidates, or both.^97–99^ In the following Foundation grant cycle, which was underway during this project, success rates were equivalent for male and female principal investigators and the policy did not need to be invoked. (See summary data in the online appendix.) In the subsequent cycle, the reviewer training module alone did not result in equivalent success rates at Stage 1 among male and female principal investigators, and the policy of equivalence was invoked.^100^

Our study had three main limitations. First, principal investigators were not randomized to one grant program or the other. Although aspects of our study minimized the potential to observe the results we found, the non-randomized design leaves open the possibility that unobserved confounders or selection bias may have contributed to observed differences. Second, our planned analyses did not include investigation of 3-way analyses between grant program group, self-reported sex, and age. It is possible that the overall two-way interaction observed might be driven by differences in evaluation of men and women who are more advanced in their careers. Our planned analyses did not include this, as we were concerned about insufficient sample size to include three-way interactions in the model. Third, we assumed that people who self-report as female or male also identify as women or men, respectively. This may not be true in all cases. Data are lacking regarding how many people identify as transgender or non-binary in Canada. Context may be offered by a recent analysis suggesting that transgender people were 0.4% of the US population.^101^ If this proportion were reflected in our study and if all transgender applicants reported their sex as their assigned sex at birth, this would represent 28 people in total within this study, a number unlikely to meaningfully change the results of our analyses.

Our study also has four main strengths. First, it was quasi-experimental. To the best of our knowledge, this is the first evidence from a study design that was not fully observational. This enables stronger conclusions than were previously possible. Second, while quasi-experimental studies have potential for selection bias,^102^ in this study, selection bias was limited by eligibility rules. Specifically, for principal investigators who had already established their careers and received funding from the CIHR, only those whose grants from traditional programs were expiring within a specific time period were permitted to apply to Foundation. This meant that a 9 portion of allocation was dependent on an external, arbitrary variable. Third, we controlled for age and domain of research, two key confounders in studies of gender bias in grant funding. Academic rank, which correlates with age, has been shown to account for gender gaps in grant funding in other studies, as has domain of research.^68,69^ Having accounted for these key confounders strengthens our findings. Fourth and finally, our study analyzed all available data over a period of five years from a major national funding agency, thus offering evidence from a large data set of real-world grant review.

### Conclusions

Bias in grant review prevents the best research from being funded. When this occurs, lines of research go unstudied, careers are damaged, and funding agencies are unable to deliver the best value for money, not only within a given funding cycle, but also long term as small differences compound into cumulative disadvantage. ‘People, not projects’ funding programs may be at higher risk of reproducing and exacerbating common societal biases in research funding. To encourage rigorous, fair peer review that results in funding the best research, we recommend that funders focus assessment on the science rather than the scientist, measure and report funding by applicant characteristics and potential confounding variables, and consider reviewer training and other policies to mitigate the effects of all forms of bias. Future research should investigate the role of confounding variables not included in our study, other types of bias beyond gender, and methods of increasing fairness and rigour in peer review.

## APPENDICES

1. Methodological and Analytical
2. Details R Code

## DECLARATIONS

### Abbreviations

CIHR: Canadian Institutes of Health Research

### Ethics Approval, Consent to Participate and Consent for Publication

The views expressed in this paper are those of the authors and do not necessarily reflect those of the CIHR or the Government of Canada. Data were held internally and analyzed by staff at the CIHR within their mandate as a national funding agency. Research and analytical studies at the CIHR fall under the Canadian Tri-Council Policy Statement 2: Ethical Conduct for Research Involving Humans (available: pre.ethics.gc.ca/eng/policy-politique/initiatives/tcps2- eptc2/Default/, accessed 2017 July 13.) This study had the objective of evaluating CIHR’s Investigator-Initiated programs, and thus fell under Article 2.5 of TCPS-2 and not within the scope of Research Ethics Board review in Canada. Nevertheless, applicants were informed through ResearchNet, in advance of peer review, that CIHR would be evaluating its own processes. All applicants provided their electronic consent; no applicant refused to provide consent.

### Availability of Data and Materials

Data are confidential due to Canadian privacy legislation. Researchers interested in addressing other research questions related to grant funding may contact the CIHR at.

### Contributions

Conceptualization: HW MH DG (see Acknowledgements); Methodology: HW JW RD AC MH DG; Formal Analysis: JW; Investigation: HW JW RD AC; Data Curation: JW RD AC; Writing – Original Draft: HW JW; Writing – Review & Editing: HW JW RD AC MH CT SS DG; Visualization: HW JW RD; Project Administration: HW RD. Contributions are listed according to the CReDIT taxonomy (docs.casrai.org/CRediT).

## Acknowledgements

The authors gratefully acknowledge early work by Ms. Anne-Sophie Julien, statistician, who worked with HW to outline potential analytical approaches, drafted preliminary R code for the proposed research question, and reviewed a draft of the manuscript. We also thank Ms. Anne Strong, CIHR staff, for extracting the dataset, and Dr. Anne Martin-Matthews, CIHR Vice-President, for her comments on drafts of this manuscript, and many colleagues who commented online in response to previous preprint versions of this manuscript. The authors are extremely grateful to Dr. Jonathan A. Whiteley (JW), Dr. Rachelle E. Desrochers (RD), Dr. Alysha Croker (AC), and Dr. Danika Goosney (DG), all of whom were employed at the CIHR during this project and assisted with the scoping of the analysis and interpretation of the results. JW conducted the analysis at the CIHR as the data used consisted of confidential applicant information that could not be shared externally. He also drafted details of the methodology in collaboration with HW. The CIHR is committed to collaborating with the research community to advance knowledge on best practices in peer review without unduly influencing the conclusions drawn by its collaborators. For this reason, CIHR employees are not currently permitted to be co-authors on papers describing analyses of the agency’s programs. Without the active participation of these CIHR employees and their contributions, this paper would not have been possible

## APPENDIX 1. Methodological and Analytical Details

### Population, Setting and Data

Beginning in 2014, the Canadian Institutes of Health Research (CIHR) phased out traditional open grant programs and divided all investigator-initiated funding into two new programs: the Project grant program and Foundation grant program. Both new programs used a staged review process in which lower-ranked applications were rejected from progressing to further stages. As in traditional programs, reviewers in the new Project grant program were instructed to primarily assess the research proposed. Seventy-five percent of the application score was based on reviewers’ assessments of ideas and methods while 25% was based on reviewers’ assessments of principal investigators’ and teams’ expertise, experience, and resources. In contrast, the Foundation grant program was about ‘people, not projects’ and was designed to provide grants to fund programs of research. At the first stage of the Foundation review process, reviewers were instructed to primarily assess the principal investigator, with 75% of the score allocated to reviewers’ assessments of principal investigators’ leadership, productivity, and the significance of their contributions, and 25% to a one-page summary of their proposed 5- or 7- year research program. Only principal investigators who passed this stage were invited to submit a detailed proposal describing their research. Thus, these new programs enabled a direct, quasi-experimental comparison of success rates of male and female applicants in grant programs with and without an explicit focus on the caliber of the principal investigator.

New investigators and those who had never held CIHR funding could apply to programs of their choice. Established principal investigators who already held CIHR funding were eligible for the Foundation program if one or more of their active CIHR grants was scheduled to end within a specific date range. Principal investigators could apply to multiple programs, with some restrictions. In the first cycle of the Foundation program, principal investigators who passed the first stage and were accepted to submit a full description of their research could not simultaneously apply to the last cycle of traditional programs. In the second cycle of the Foundation program, principal investigators could apply to Foundation and Project programs, providing they did not submit the same research proposal to both programs.

We analyzed data from all applications submitted to CIHR grant programs across all investigator-initiated competitions in 2011 through 2016. We excluded applications that were withdrawn, as these did not receive full peer review. We also excluded applications if the principal investigator, referred to as the nominated principal applicant in the CIHR system, had not reported their sex, their age, the domain of research of their application, or if their self-reported age was unrealistic. We defined unrealistic ages as occurring when a principal investigator’s self-reported birth year was prior to 1920 or after 2000. Ensuring correct date entry in web-based forms is a known challenge in human-computer interaction.^1^ In the online system used to collect personal data in this study, the Canadian Common CV, the default birth date is prior to 1920, suggesting that such self-reported birth years were most likely to occur when people did not enter a birth date. Birth years in the 2000s were deemed to be errors and may have occurred, for example, when people accidentally entered the current year as they created or updated their Canadian Common CV rather than their birth year.

### Data Source

Our dataset was a full export of CIHR competition data from its Electronic Information System (EIS). This dataset does not include withdrawn applications. It only includes applications submitted that were fully assessed by peer review and either approved or not.

### Analysis

The binary outcome of interest was application success. In the model, we coded this as true (1) if the application was approved after the peer review process, and false (0) if not. Because our aim was to analyze the effects of peer review, we coded applications that were deemed fundable but not approved in the competition to which they were applied as unsuccessful, even if they were later awarded money through other administrative processes such as bridge grants or priority announcements for specific funding areas.

We modeled grant success rates as a function of the grant program, principal investigators’ self-reported binary sex (male or female), self-reported age, self-declared domain of research (biomedical; clinical; health systems and services; or social, cultural, environmental and population health),^2^ and an interaction term between each principal investigator’s sex and the grant program to which they were applying. Grant programs were grouped into three categories: traditional investigator-initiated grant programs programs (includes regular open operating grant programs from fiscal years 2011/12 to 2014/15 which account for 88% of grants in the Traditional programs group plus 6 smaller programs, some of which continued into fiscal year 2015/16); Project (spring 2015 competition, fiscal year 2015/16); and Foundation (two competitions: 2014/15 and 2015/16). All predictors were categorical variables, except for applicant age, which was continuous and was mean-centered prior to analysis.

The interaction term allowed us to address the objective of this study by determining whether there was any effect of different review criteria on relative success rates between male and female applicants after controlling for age and domain of research. We controlled for these because younger cohorts of investigators included larger proportions of female principal investigators, as did domains of health research other than biomedical. The adjustment for age also helped account for the fact that both the Foundation program and Project program had a predefined minimal allocation to new investigators, meaning those within their first five years as independent investigators. The CIHR collected data about binary sex, not gender; therefore, our study assumes that people who self-reported as female or male identified as women or men, respectively. The CIHR did not collect complete data on other applicant characteristics that have been shown to be associated with disparities in funding and career progression; for example, career stage, race, ethnicity, Indigeneity and disability.^3,4^

We used Generalized Estimating Equations (GEE) to fit a logistic model that accounted for the same principal investigator submitting multiple applications,^5,6^ including principal investigators who applied to both Project and Foundation programs. We conducted analyses in R statistical computing software, version 3.4.0,^7^ using the geepack package to fit models.^8^ We then used the fitted model to test the pairwise effect of sex within each program, using the lsmeans package.^9^ This allowed us to compute marginal effects for specific contrasts of interest.

The R code used for our analyses is appended, with variable names masked for security and privacy reasons.

### Results

There were a total of 25,706 applications during the five years of this study. We excluded 1,788 applications consisting primarily of principal investigators with unrealistic years of birth; i.e., birth years prior to 1920 (n=1,631) or after 2000 (n=12). The final dataset analyzed contained 23,918 applications from 7,093 unique principal investigators. There were 15,775 applications from 4,472 male principal investigators and 8,143 applications from 2,621 female principal investigators. Twenty-eight percent of principal investigators submitted a single application during the five-year study period, 20% submitted two applications, 25% submitted three or four applications, and the remaining 27% of principal investigators submitted five or more applications. The maximum number of applications from a principal investigator during the 5- year period was 40.

The overall grant success rate across the data set was 15.8%. After adjusting for age and research domain, the predicted probability of funding was 0.9 percentage points higher for male principal investigators than female principal investigators in traditional programs (OR 0.934, 95% CI 0.854-1.022). This gap was 0.9 percentage points in the Project program (OR 0.998, 95% CI 0.794-1.229) and 4.0 percentage points in the Foundation program (OR 0.705, 95% CI 0.519-0.960). In other words, female applicants experienced significantly lower success rates than male applicants in Foundation, but not in Project nor in traditional programs. This was confirmed by using the model coefficients to compute contrasts and associated odds ratios for sexes within each program. Across all grant programs, odds of receiving funding were also lower in the three non-biomedical research domains and for younger principal investigators.

Tables S1 and Table S2 provide different ways of viewing these results. Table S1 shows the raw GEE results while Table S2 shows the observed success rates, predicted probabilities, and calculated odds ratios for the interaction between applicant sex and program at uniform values of age, averaged across domains.^9^ Table S3 shows results by review stage for the cycles included in the quasi-experiment and thus in the analyses in the paper. Table S4 shows results by review stage for the following cycle after changes were made to the program, specifically, after a reviewer learning module on the topic of unconscious bias was implemented in the program.

## TABLES

**Table S1.** Odds of Grant Success: GEE Results

|  | Odds Ratio (OR) | OR lower bound of 95% confidence interval | OR upper bound of 95% confidence interval |
| --- | --- | --- | --- |
| (Intercept): Male applicants in Program: traditional programs, Research Domain: Biomedical, Mean age | 0.229 | 0.216 | 0.242 |
| Female applicants | 0.934 | 0.854 | 1.022 |
| Program: Project | 0.762 | 0.675 | 0.860 |
| Program: Foundation | 0.748 | 0.641 | 0.873 |
| Age (mean-centered) | 1.005 | 1.001 | 1.010 |
| Research Domain: Clinical | 0.815 | 0.738 | 0.900 |
| Research Domain: Health Systems and Services | 0.846 | 0.747 | 0.959 |
| Research Domain: Population and Public Health | 0.772 | 0.681 | 0.877 |
| Sex * Program: Female applicants in Project | 0.988 | 0.794 | 1.229 |
| Sex * Program: Female applicants in Foundation | 0.705 | 0.518 | 0.960 |

**Table S2.** Associations between Predictors and Funding Success

|  | Number of applications in data set | Percent successful (unadjusted) | Predicted probabilities* (after adjusting for age and research domain) | Odds Ratio (OR)* | OR lower bound of 95% confidence interval* | OR upper bound of 95% confidence interval* |
| --- | --- | --- | --- | --- | --- | --- |
| **Applicant Sex x Program Interaction** |  |  |  |  |  |  |
| Male applicants in traditional programs (reference) | 11879 | 17.3% | 16.3% | 1.000 |  |  |
| Female applicants in traditional programs | 6326 | 15.7% | 15.4% | 0.934 | 0.854 | 1.022 |
| Male applicants in Project | 2469 | 13.5% | 12.9% | 0.762 | 0.675 | 0.860 |
| Female applicants in Project | 1119 | 12.0% | 12.1% | 0.703 | 0.587 | 0.841 |
| Male applicants in Foundation | 1427 | 13.9% | 12.7% | 0.748 | 0.641 | 0.873 |
| Female applicants in Foundation | 698 | 9.2% | 8.8% | 0.493 | 0.375 | 0.647 |
| **Research Domain** |  |  |  |  |  |  |
| Biomedical (reference) | 14159 | 16.9% | n/a | 1.000 |  |  |
| Clinical | 4497 | 14.3% | n/a | 0.815 | 0.738 | 0.900 |
| Health Systems and Services | 2609 | 14.9% | n/a | 0.846 | 0.747 | 0.959 |
| Population and Public Health | 2653 | 13.6% | n/a | 0.772 | 0.681 | 0.877 |
| **Age** |  |  |  |  |  |  |
| Age (mean-centered) | 23918 | 15.8% | n/a | 1.005 | 1.001 | 1.010 |

\*Predicted probabilities and odds ratios (including 95% confidence intervals) calculated with lsmeans package.

**Table S3.** Foundation results by review stage during study

| **Cycle** | **Stage** | **Male applicants:**<br>**n (% of previous**<br>**stages)** | **Female applicants:**<br>**n (% of previous**<br>**stages)** |
| --- | --- | --- | --- |
| **2014/15** | Applied at Stage 1 | 850 | 490 |
| *Review focus: caliber*<br>*of applicant* | Invited to Stage 2<br>and applied | 317<br>(317/850 = 37%) | 126<br>(126/490 = 26%) |
| *Review focus: quality*<br>*of research* | Funded | 109<br>(109/317 = 34%,<br>109/850 = 13%) | 41<br>(41/126 = 33%,<br>41/490 = 8%) |
| **2015/16** | Applied at Stage 1 | 623 | 279 |
| *Review focus: caliber*<br>*of applicant* | Invited to Stage 2<br>and applied | 198<br>(198/623 = 32%) | 62<br>(62/279 = 22%) |
| *Review focus: quality*<br>*of research* | Funded | 91<br>(91/198 = 46%,<br>91/623 = 15%) | 29<br>(29/62 = 47%,<br>29/279 = 10%) |

A subset of applications also underwent additional Stage 3 review after Stage 2, again with a focus on the quality of the research, before being funded or rejected. There were no differences by applicant's self-reported sex.

**Table S4.** Foundation results by review stage after implementation of unconscious bias reviewer training module

| **Cycle** | **Stage** | **Male applicants:**<br>**n (% of previous**<br>**stages)** | **Female applicants:**<br>**n (% of previous**<br>**stages)** |
| --- | --- | --- | --- |
| **2016/17** | Applied at Stage 1 | 428 | 172 |
| *Review focus: caliber*<br>*of applicant* | Invited to Stage 2<br>and applied | 165<br>(165/428 = 39%) | 69<br>(69/172 = 40%) |
| *Review focus: quality*<br>*of research* | Funded | 54<br>(54/165 = 33%,<br>54/428 = 13%) | 22<br>(22/69 = 32%,<br>22/172 = 13%) |

## APPENDIX 2. R Code

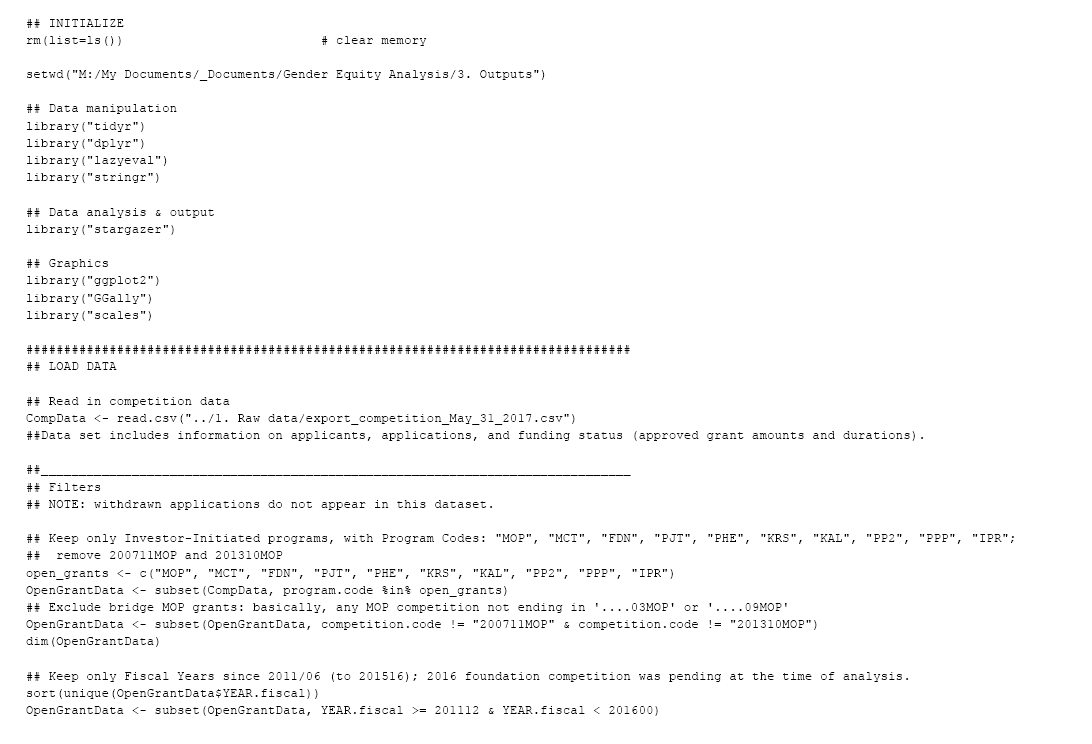
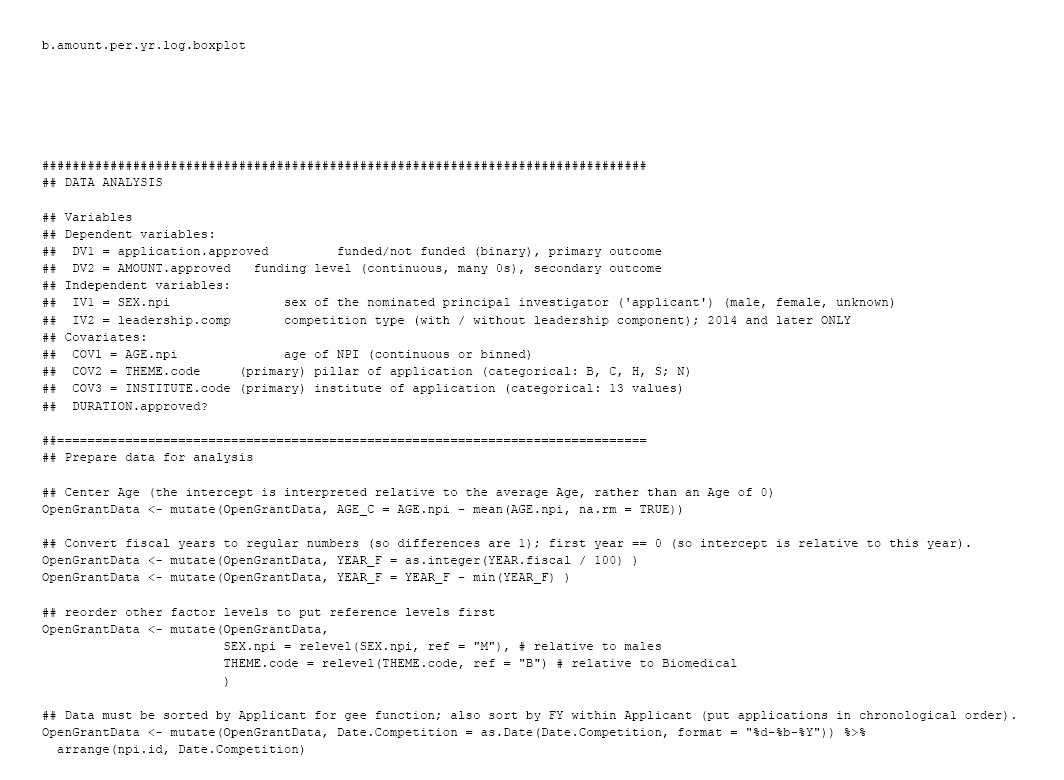
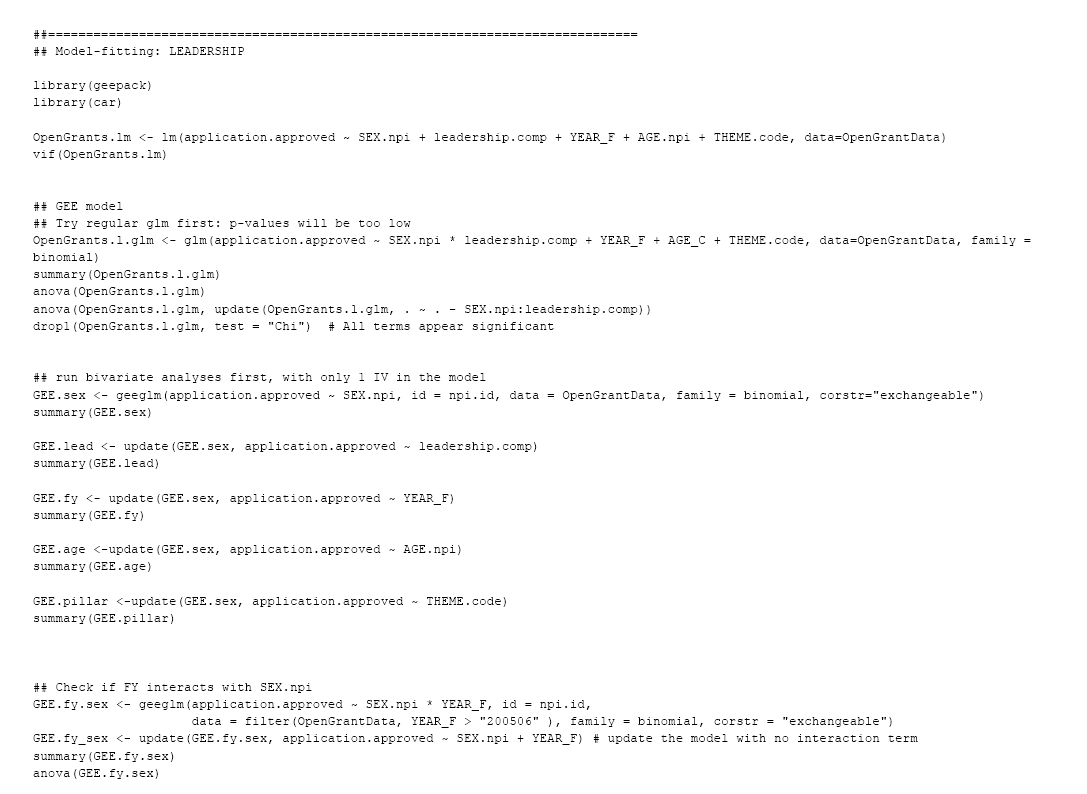
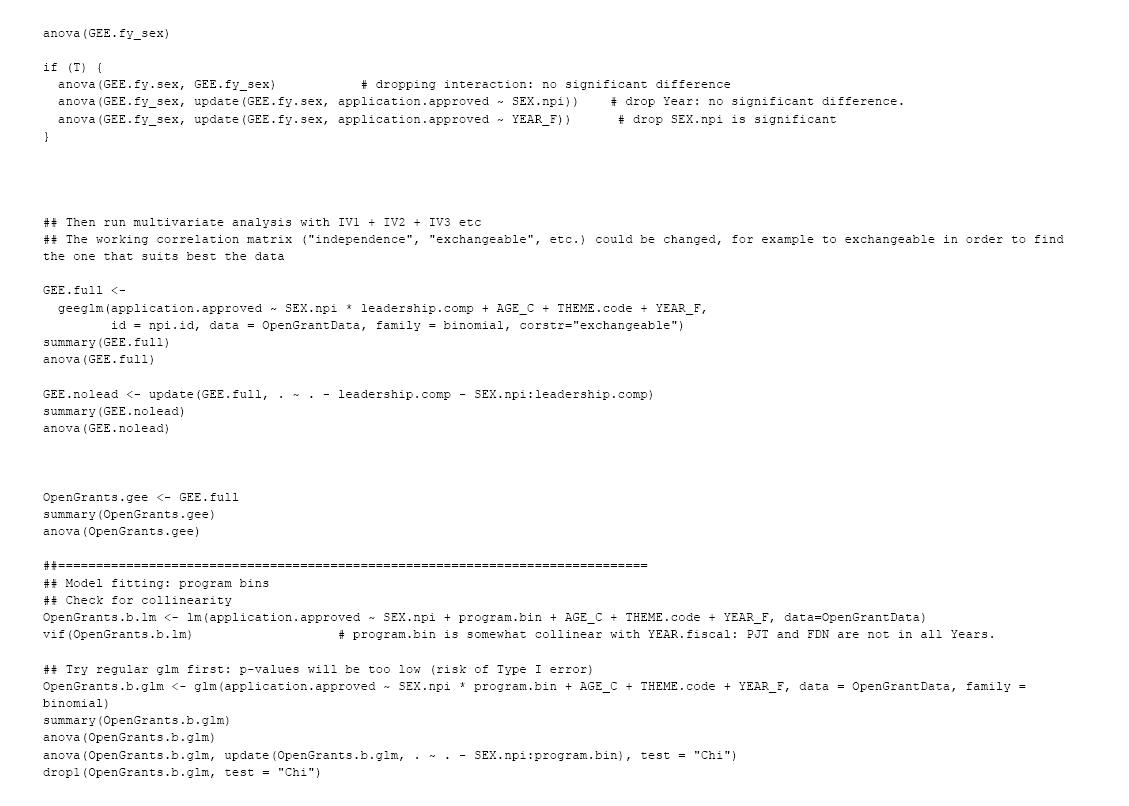
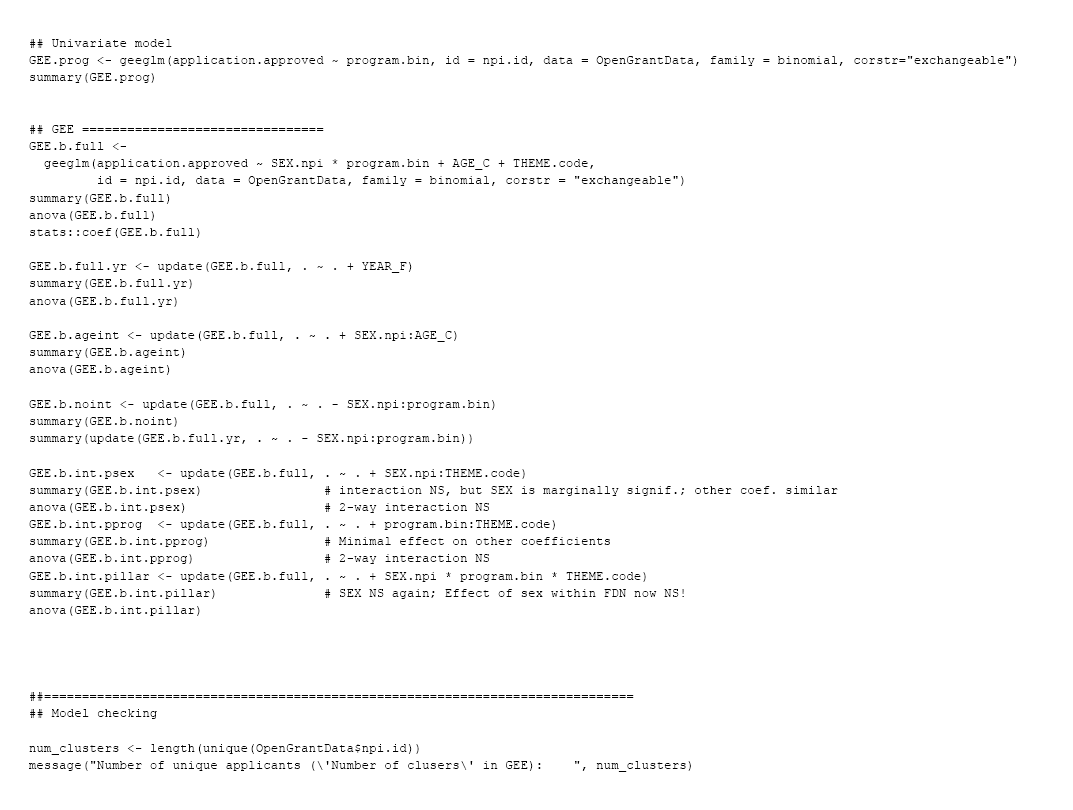
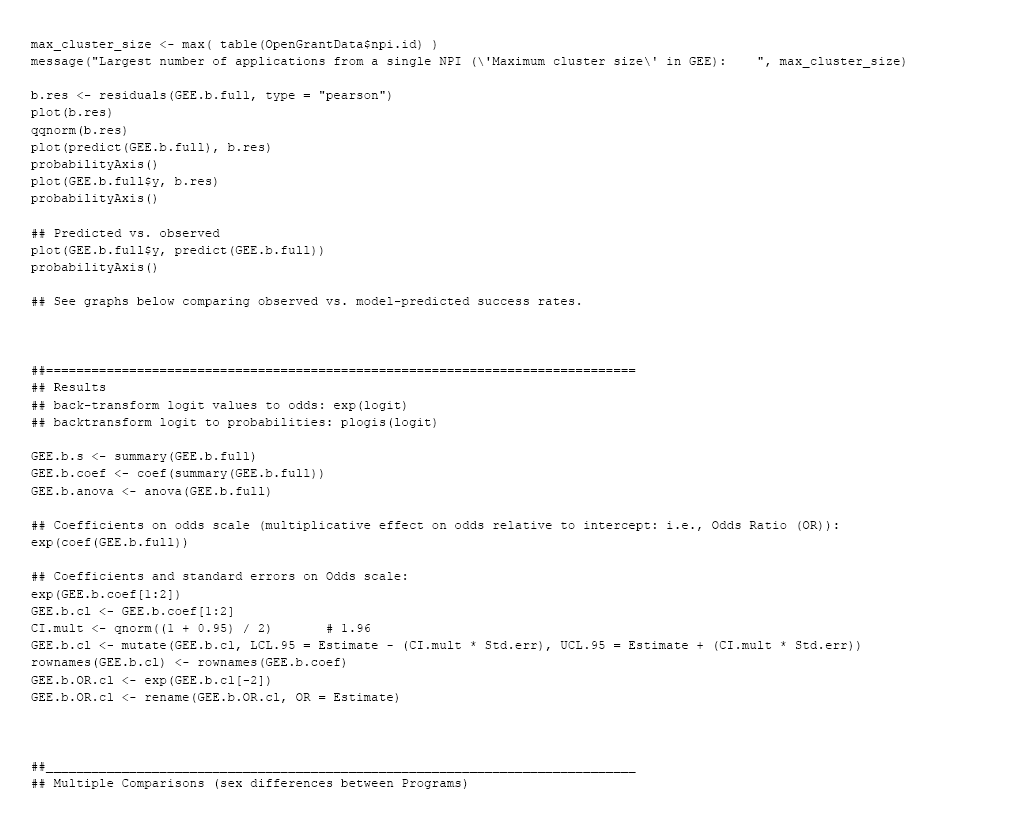
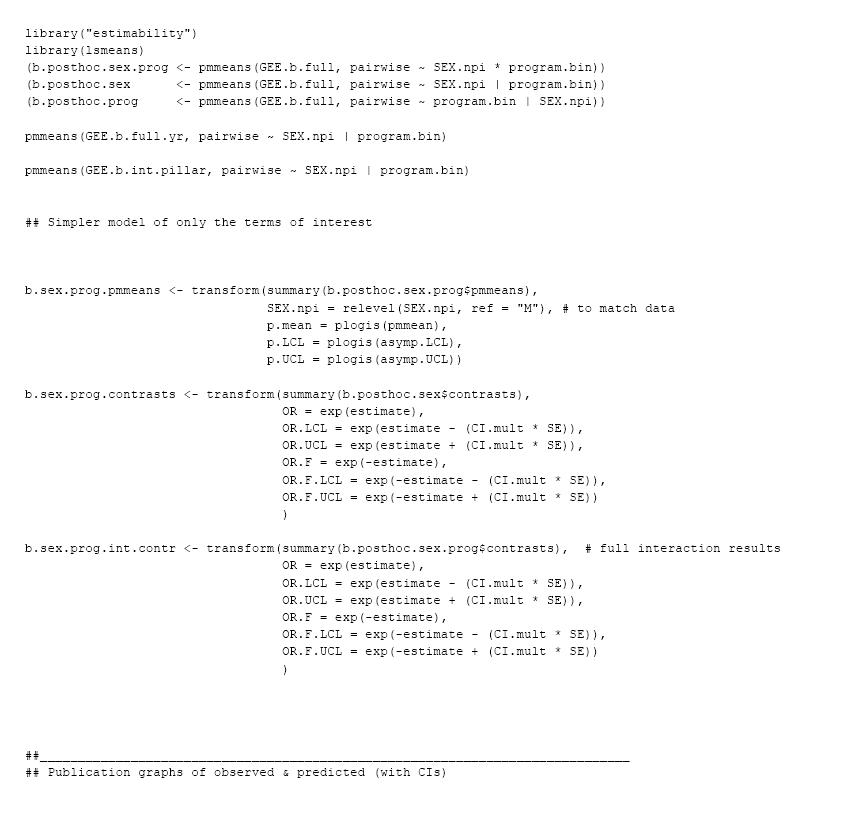
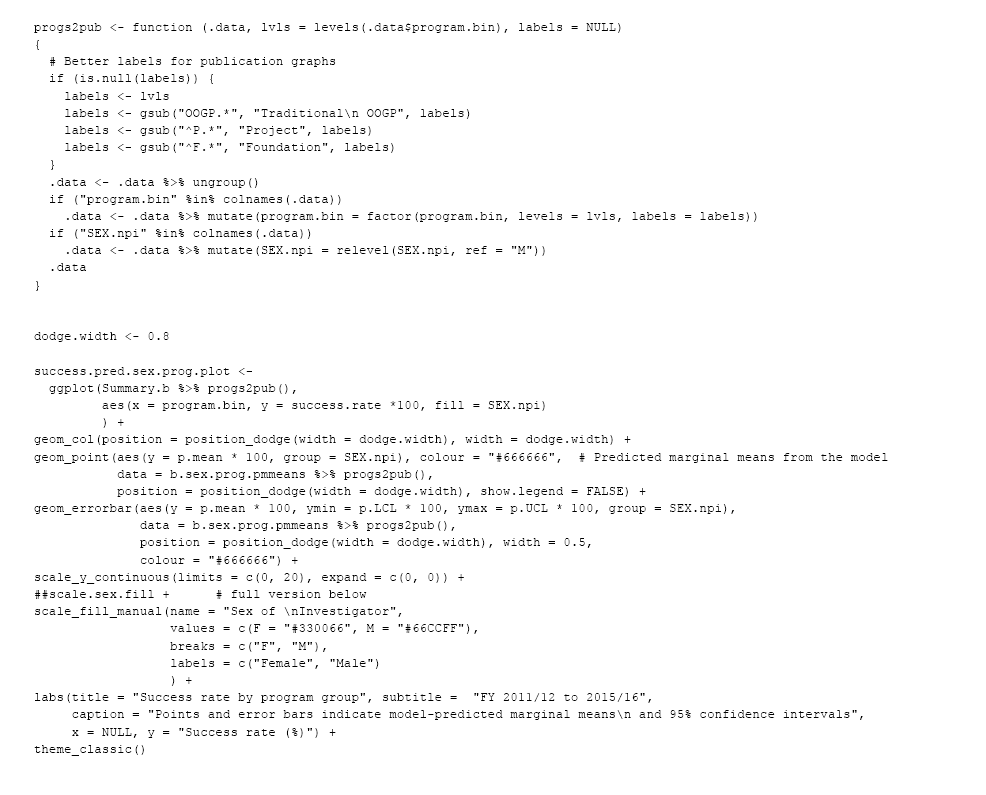
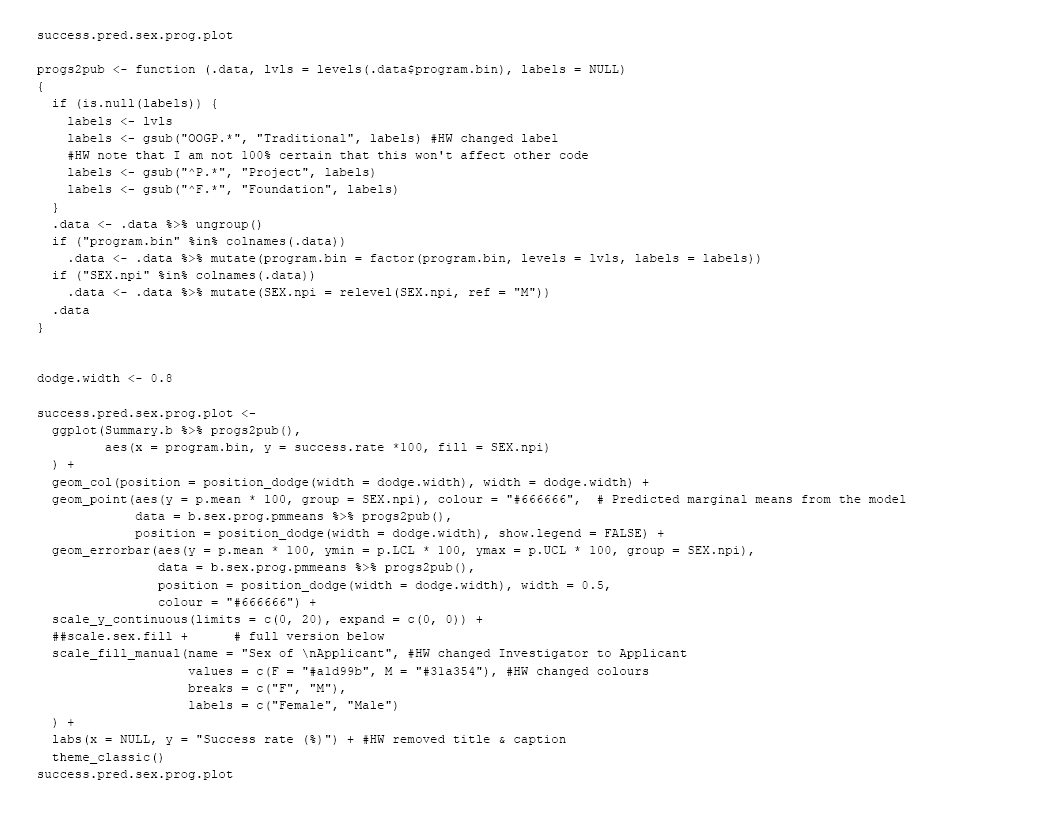
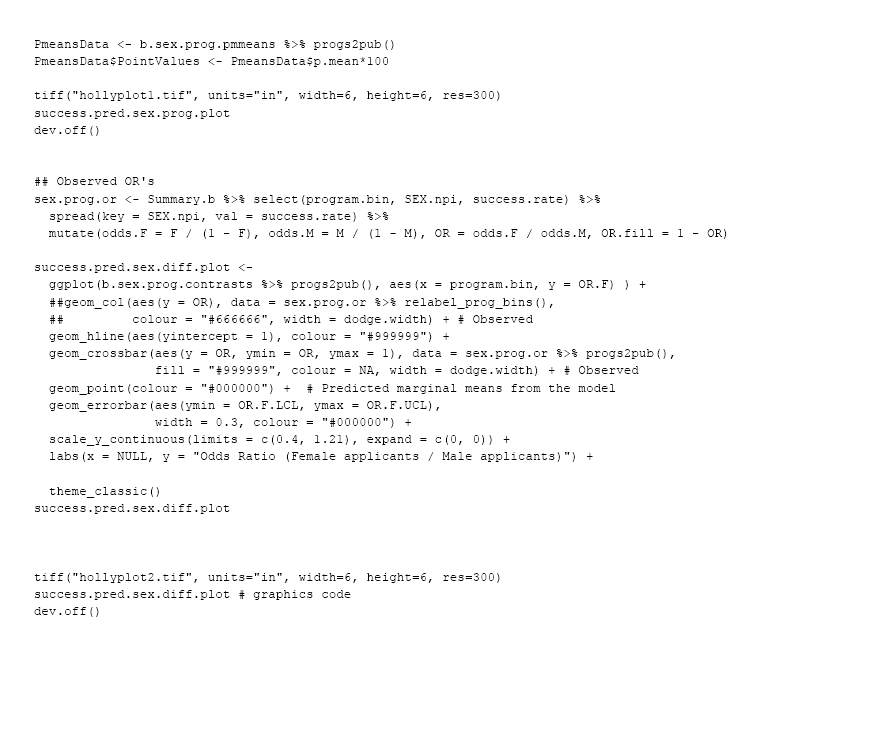
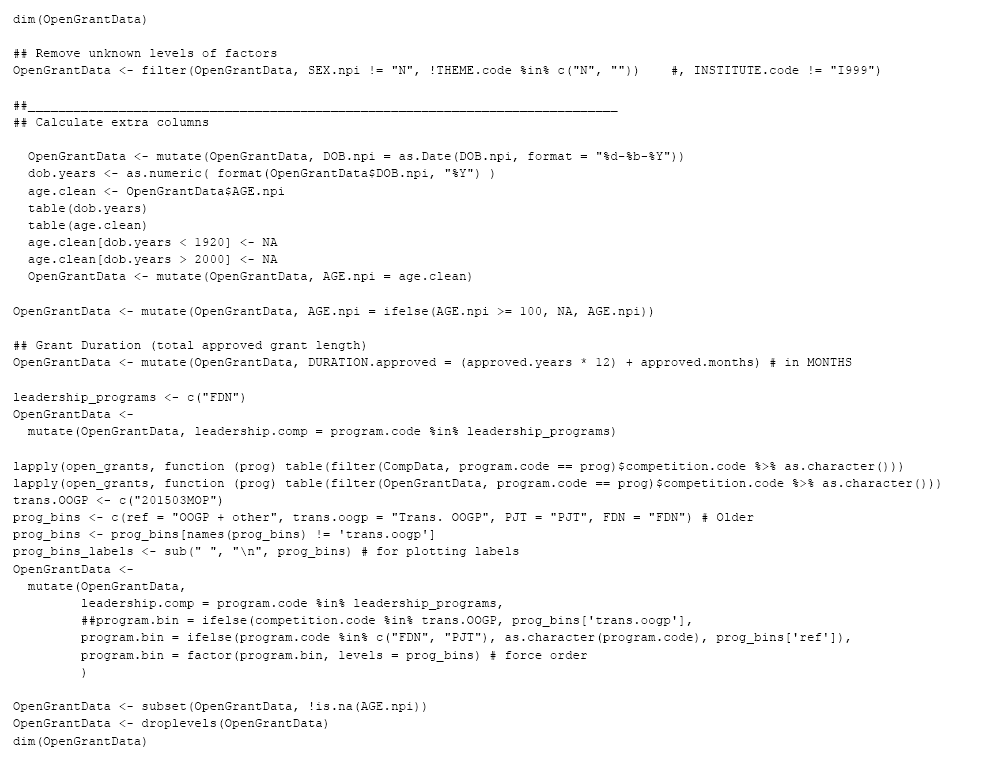
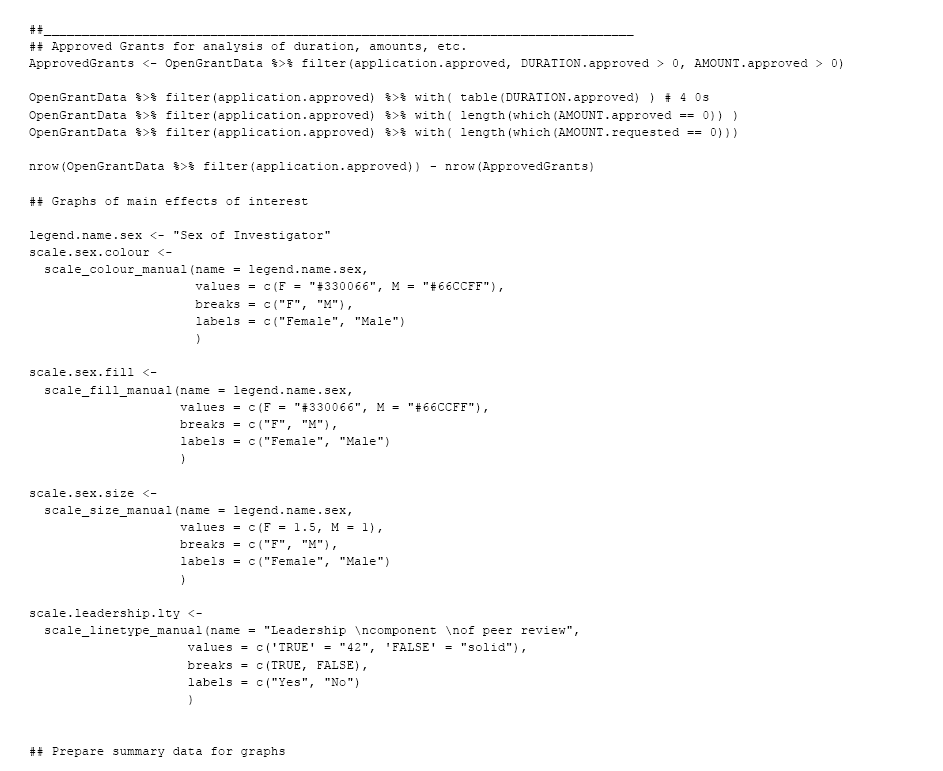
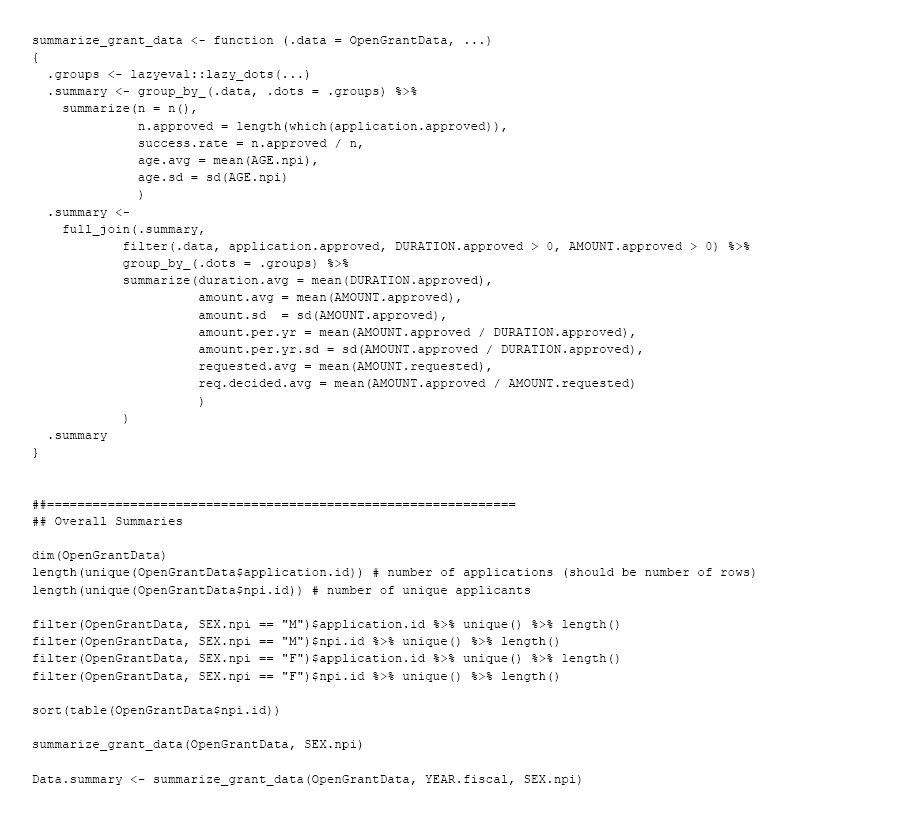
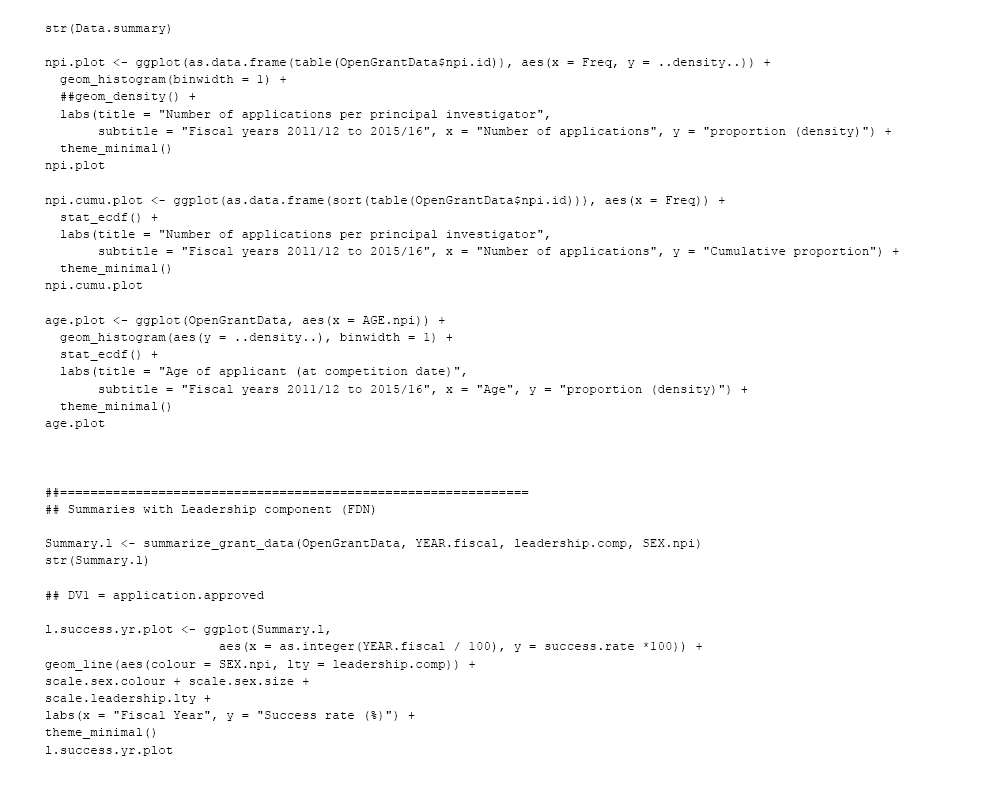
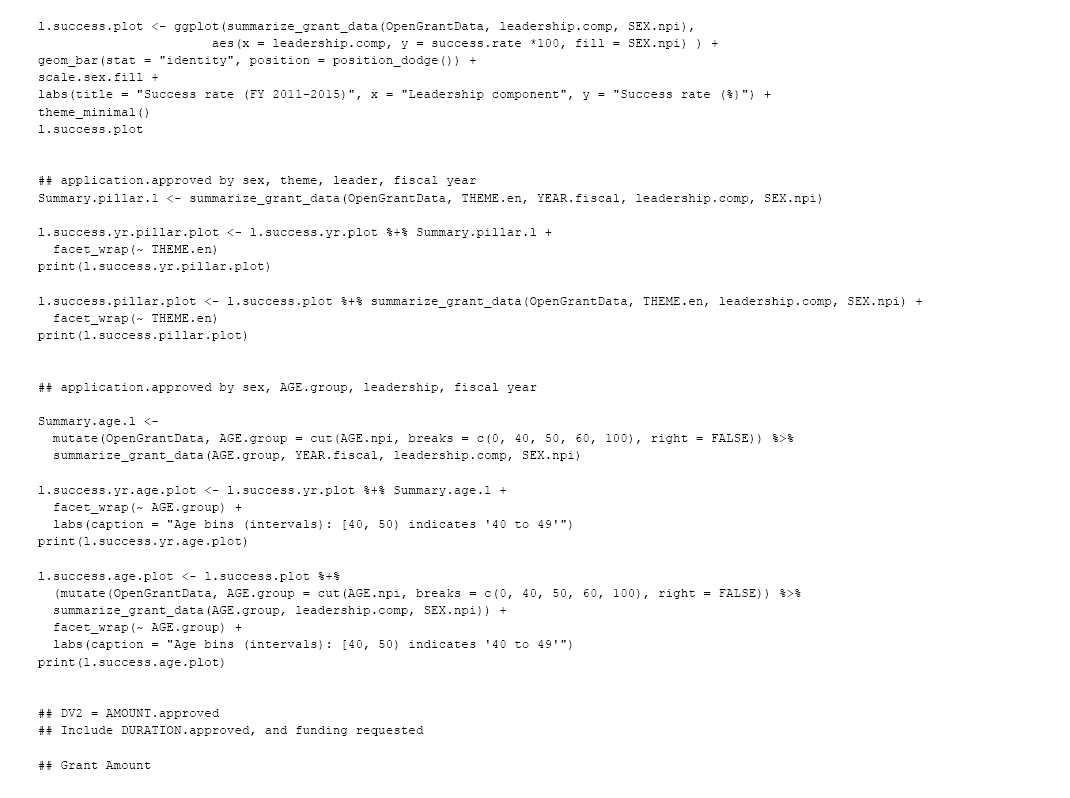
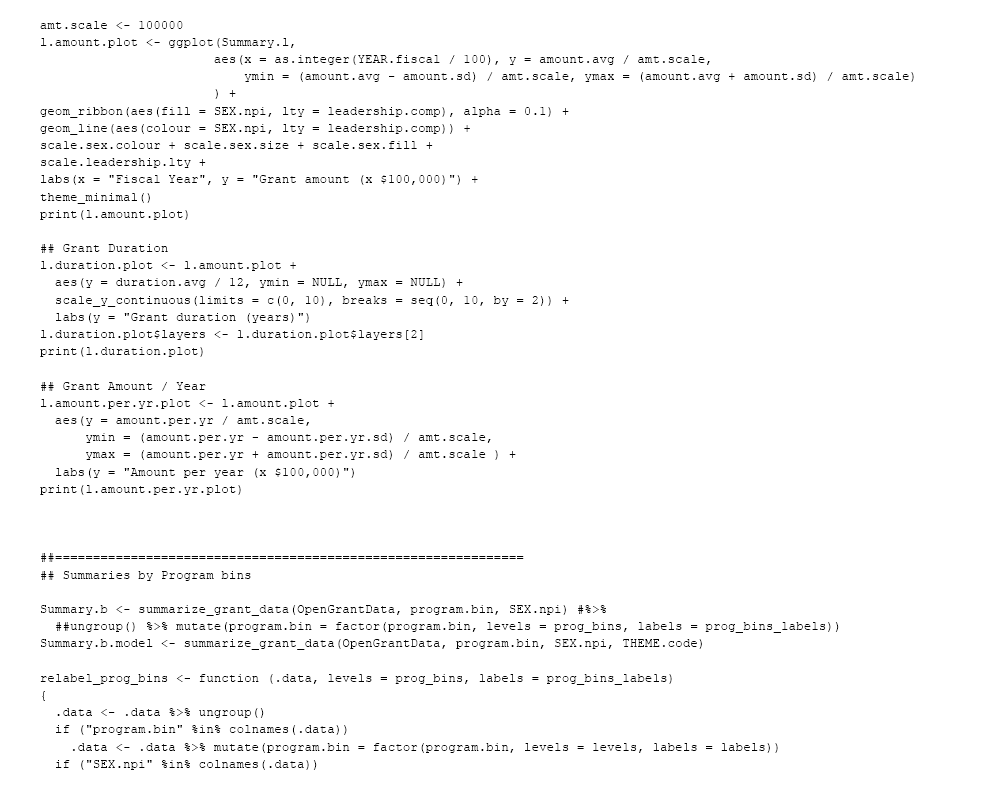
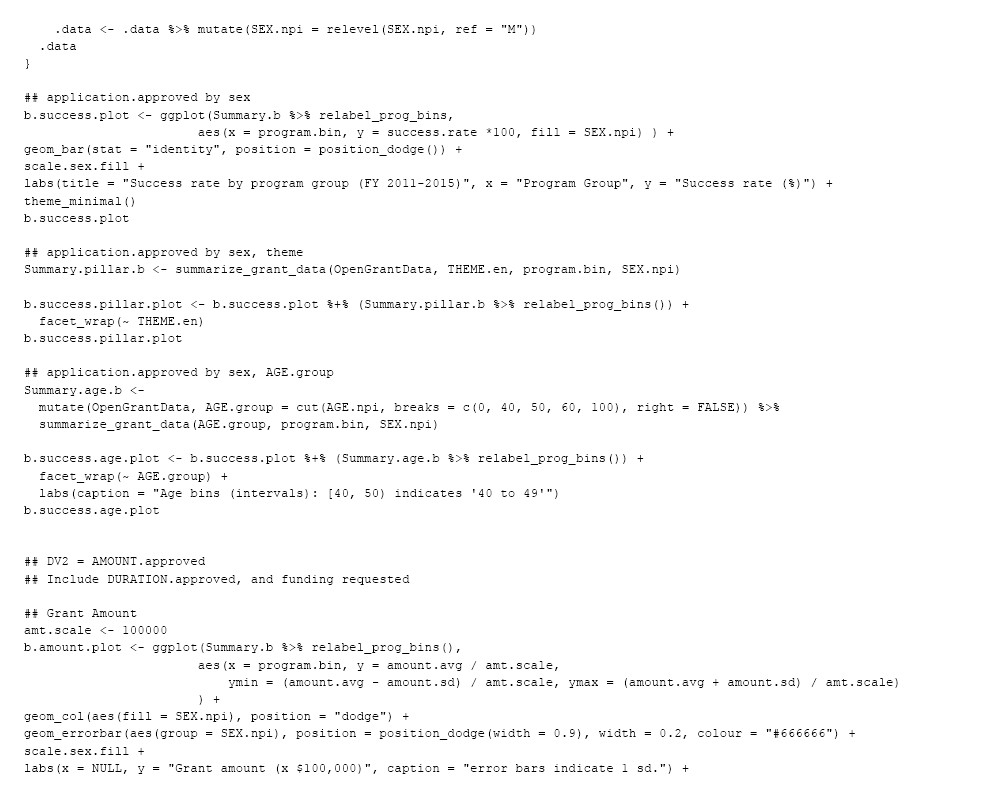
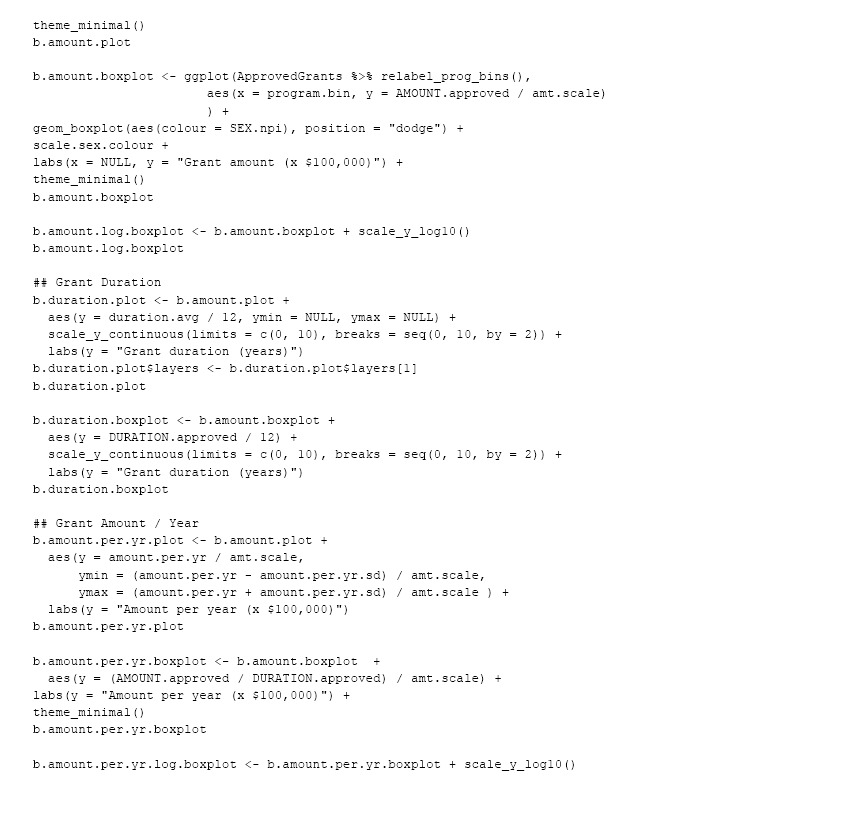

